# A high-resolution genomic study of the Pama-Nyungan speaking Yolngu people of northeast Arnhem Land, Australia

**DOI:** 10.1101/2025.04.03.646226

**Authors:** Neville White, Manoharan Kumar, David Lambert

**Author notes:** **Correspondence:** Neville White, Department of Environment and Genetics, La Trobe University, Melbourne, Victoria 3086, Australia. **Funding information:** We thank the Australian Research Council for the following grants: DP170101313 and LP120200144.

## Abstract

**Objectives:** About 300 Aboriginal languages were spoken in Australia. These were classified into two groups: Pama-Nyungan (PN), comprised of one language Family, and Non-Pama-Nyungan (NPN) with more than 20 language Families. The Yolngu people belong to the larger PN Family and live in Arnhem Land in northern Australia. They are surrounded by groups who speak NPN languages. This study, using nuclear genomic and mitochondrial DNA data, was undertaken to shed light on the origins of the Yolngu people and their language. The nuclear genomic sequences of Yolngu people were compared to those of other Indigenous Australians, as well as Papuan, African, East Asian and European people.

**Materials and methods:** With the agreement of Indigenous participants, samples were collected from 13 Yolngu individuals and 4 people from neighbouring NPN speakers and their nuclear genomes sequenced to a 30lil coverage. Using the short-read DNA BGISEQ-500 technology, these sequences were mapped to a reference genome and identified ∼24.86 million Single Nucleotide Variants (SNVs). The Yolngu SNVs were then compared to those of 36 individuals from 10 other Indigenous populations/locations across Australia and four worldwide populations using multidimensional scaling, population structure, F3 statistics and phylogenetic analyses.

**Results:** Using the above methods, we infer that Yolngu speakers are closely related to neighbouring NPN speakers, followed by the Weipa population. No European or East Asian admixture was detected in the genomes of the Yolngu speakers studied here, which contrasts with the genomes of many other PN speakers that have been studied. Our results show that Yolngu speakers are more closely related to other PN speakers in the northeast of Australia than to those in central and western Australia studied here. Yolngu and the other Australian populations from this study share Papuans as an out-group.

**Discussion:** The study presented here provides an account of the nuclear and mitochondrial genomic diversity within the PN Yolngu Aboriginal population. The results show the Yolngu sample and their NPN neighbours have a strong genetic relationship. They also offer evidence of ancestral links between the Yolngu and PN-speaking populations in Cape York. From earlier fingerprint studies, consistent with the genomic results shown here, we suggest that there was a movement of people from the east into northeast Arnhem Land, associated with the flooding of the Sahul Shelf, and that this occurred between about 11 Kya and 8 Kya ago. Several Yolngu myths point to such a movement. It is suggested that the spread of the PN language or its speakers may have influenced the population structure of the Yolngu. Further genomic studies, with larger samples, of populations to the east of the Yolngu around the Gulf of Carpentaria into Cape York are required to test this hypothesis. Our results imply that PN did not spread with the movement of people across the continent, rather, the PN languages diffused among the different populations. It seems clear that the languages dispersed and not the people. The low level of relatedness detected between the Yolngu people and the people of the central arid desert of Australia suggests a long period of separation with different patterns of migration. Beyond Australia, Yolngu are most closely related to the Papuan people of New Guinea.

## 1 Introduction

Although still debated, the archaeological and dating evidence suggests that Aboriginal people arrived in northern Australia / Sahul ∼65 Kya (thousand years ago) (Clarkson et al., 2017; Allen & O’Connell, 2020).

However, it is generally accepted that after the initial colonization event(s), modern humans migrated to the arid centre of the continent by ∼50 Kya. This date is supported by a large amount of evidence (see McDonald et al. 2018 for a review). Additional support for this chronology has come from the study of mitochondrial sequence variation (Tobler et al., 2017).

“There is a substantial body of gene frequency data (mostly from single gene blood systems) compiled from Aboriginal Australians. These are best summarised in Kirk (1965), Balakrishnan et al. (1975) and Cavalli-Sforza et al. (1994)” (White, 1997, p.52). However, it was not until Rasmussen et al. (2011) that the first whole genome study was reported, based on an Aboriginal Australian man from Kalgoorlie in southwestern Australia. This genome was recovered from a 100-year-old hair sample. Rasmussen et al. (2011) suggested that Aboriginal Australians are descendants of an early modern human dispersal into East Asia approximately 62-75 Kya. Later, Malaspinas et al. (2016) conducted a comprehensive study of 83 genomes of Aboriginal Australians from eight populations, in addition to 25 Papuan genomes. The authors provided evidence suggesting that Aboriginal Australians were part of a single Out-of-Africa event and estimated the divergence time between Eurasians and Australo-Papuans at 51-72 Kya. Aboriginal Australians are estimated to have diverged from Papuans 47 Kya (Silcocks et al., 2023). Recent whole genome studies of a large number of Aboriginal Australian populations have enabled an improved estimate both of the timing of the arrival and of the subsequent migration events within the continent (Allen et al., 2020).

In parallel with these genetic studies, there has been considerable interest in the origins of the languages spoken by Aboriginal Australians (e.g. Walsh, 1991; Evans, 2020; Bowern, 2023). At the time of first European settlement, it is estimated that Aboriginal Australians spoke about 300 different languages. The majority of these belonged to the PN language Family, the speakers of which cover 85% of the Australian continent (Wurm, 1971). The remaining 15% is occupied by NPN speakers, with more than 20 language Families (Evans, 2020), whose distribution ranges from the Gulf of Carpentaria, through Arnhem Land to the Kimberley (Capell, 1940; 1942; O’Grady et al., 1966; Wurm, 1971). The region occupied by NPN speakers has the greatest linguistic diversity in the continent (Wurm, 1971; Evans, 2020) (Figure 1). This division between PN and NPN has been argued to have occurred within the Australian continent after initial colonization (Capell, 1962; Tryon, 1971; Evans, 2020). Capell (1956; 1962; 1967) pointed to the essential relatedness of Australian languages, although regional differences in both structure and vocabulary do occur. Capell (1962) considered that Australian languages cannot be linked outside Australia and that it is necessary to postulate a lengthy (perhaps 12-15 Kya) occupation of the continent to account for the present diversity among language groups. Capell (1956) argued that the NPN languages did not originate in their present geographic locations and have no obvious relationship to languages beyond Australia. Heath (1990) and Evans (2020) suggested the possibility of a common origin of Aboriginal Australian languages using phonetic similarities. However, this suggestion is still debated (Bowern, 2010). Bouckaert et al. (2018) regard PN languages as having a relatively recent origin, estimated at being between 6 Kya and 13 Kya.

**FIGURE 1:**
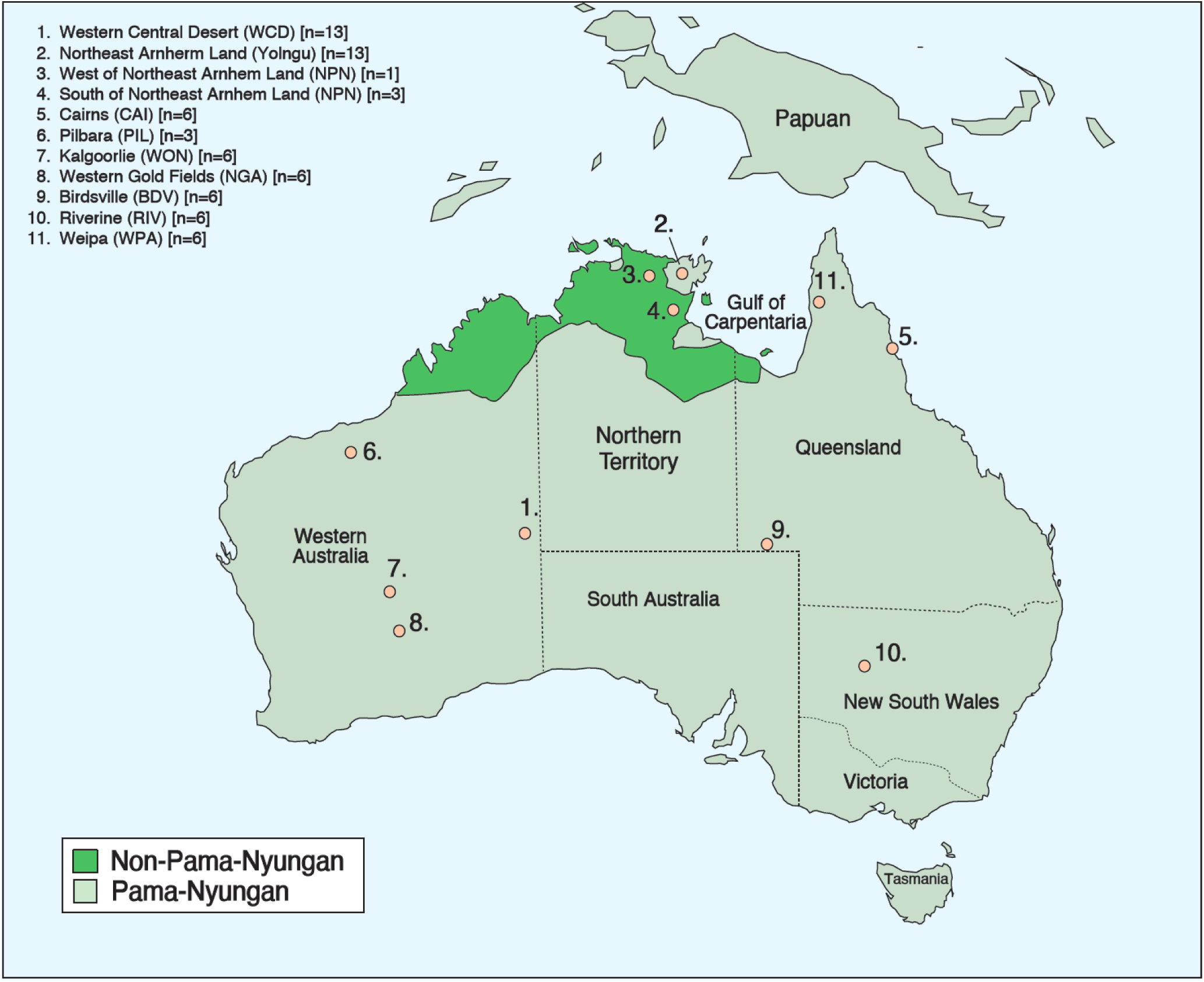
Geographic locations of the 11 Aboriginal Australian populations (1-11) and their language Family (highlighted in yellow) sequenced as part of this study. The distributions of PN (light green) and NPN (dark green) languages are shown.

The Yolngu are an enclave of PN-speaking people in northeast Arnhem Land within a band of NPN-speaking people (Figure 1). They are also referred to in the early literature as Murngin, Wulamba, and Malag, and have been extensively studied. Much of this literature relates to the Yolngu’s social organization, religion, art and rituals (Warner 1958; Thomson 1949a,b; Berndt 1951, 1952; Williams 1986; Morphy 1991; White 1995). The Yolngu language differs in both structure and vocabulary from all of the NPN languages that surround them. The language is differentiated into a number of exogamous dialects, some of which differ to such an extent that they are almost mutually unintelligible.

The Yolngu have also been the subject of several early quantitative genetic studies (White, 1979, 1995; Parsons & White, 1976). These studies were conducted on fingerprint total ridge counts, a trait which is determined by a number of additive genes with a high heritability (the highest of any metric trait recorded). These studies found that, as a group, the Yolngu are genetically distinct from neighbouring language units, although the westernmost dialects are not strongly differentiated from their immediate neighbours, the Rembarrnga. Nor are the Yolngu southern and western dialects differentiated from their eastern neighbours, the Anindilyakwa on Groote Eylandt. There was shown to be a close similarity in the total ridge count of Yolngu people as a whole and PN-speaking people to the east around the Gulf of Carpentaria, namely the Yanyuwa and Garawa people, and those from Mornington Island, who speak an NPN language. This, together with other genetic data based on blood groups, as well as cultural information, led White (1997) to suggest that the Yolngu had their origins to the east of their present location. On the other hand, there was a particularly strong genetic differentiation based on fingerprints between the PN-Yolngu and PN-speaking groups from Central Australia (White 1997).

Until the recent articles by Reis et al. (2023) and Silcocks et al. (2023), which include a sample of NPN-speaking Tiwi people, published accounts of genomic variation among Indigenous Australians were confined to PN speakers (e.g. Malaspinas et al., 2016). The studies by Reis et al. (2023) and Silcocks et al. (2023) also included a sample of Yolngu from Galiwin’ku on Elcho Island in northeast Arnhem Land. They presented detailed accounts of genomic structural variation across four remote Aboriginal communities in northern and central Australia. In our study, we focus on the population genetic relationships and pre-history, particularly of the Yolngu people with nine other Indigenous Australian populations. Furthermore, there have been no suggestions about the role that the Yolngu population played in the settlement of Australia. This article reports the results of a study in which complete nuclear genomes of 13 Yolngu speakers from northeast Arnhem Land were sequenced and compared with genomes of people from Western Central Desert and some other PN-speaking Indigenous groups from throughout the continent.

## 2 MATERIALS AND METHODS

### 2.1 Background to the study

DNA from Yolngu people living in northeast Arnhem Land and linked to the Gapuwiyak community were analysed in this study. White obtained the saliva samples with the signed written consent of the participants and with the approval for use of their DNA in relation to the Griffith University Human Ethics Committee Approval 2022/207. The process of consultation was consistent with CARE principles. Letters of support from the community were written by the Council Secretary/Assistant Teacher, who is highly regarded and trusted in the community. This person also has excellent literacy skills in English and Djambarrpuyngu (the local dialect) and was a great help with the collection of specimens and support for the project. The tradition-oriented Yolngu community did not want to have the participants or community identified in this publication, a request which we respect throughout the manuscript. Permission to work with the DNA collected in this study was stipulated as follows: “DNA was collected from us by Neville White, who has lived and worked with us for a very long time. We ask that the DNA still belong to us […] We do not want the DNA to be given to anyone else without our permission”.

This project was explained in the Djambarrpuyngu dialect orally, given that no other adult in the community is literate in either English or their own languages. The research questions were explained at some length. An account of this, written in Djambarrpuyngu and English, was given to the Griffith University Human Research Ethics Review (Full Research Ethics Clearance was provided in April 2022). The research questions were developed in a meeting with the homeland residents, who were most interested in recording their history to use in the community school and for the development and management of their cultural landscape with other Yolngu clans and neighbouring tribes. The Council Secretary wrote, “we agree to give some spit, to use in our work together, looking at where Yolngu people came from a long time ago, following the pathways of our ancestors and relationships with other Aboriginal people. […] When the work is finished, we will return what is left to you or wash it away. It’s up to you.” It must be understood that scientific aspects of the study are not easily translatable. Relevant documents together with signed informed consent forms are held by Prof Lambert, in accordance with the requirements of the Griffith University Human Ethics Committee.

### 2.2 Sample DNA extraction and Whole Genome sequencing

We sequenced whole genomes of 17 Aboriginal Australians from northeast Arnhem Land (Figure 1) which comprised 13 Yolngu and 4 NPN speakers. We used Oragene DNA saliva kits to collect saliva. For comparative purposes we also compared these sequences with 81 genomes of Aboriginal Australians we collected from other regions of Queensland and from throughout the continent (see 2.3 Bioinformatics Analyses section below).

#### 2.2.1 DNA Extraction

A portion of the saliva sample (400 µL) was pipetted into a fresh 2 mL Eppendorf tube. The solution was incubated (56°C, 60 min) with shaking, then ammonium acetate (10 M, 150 µL) was added and mixed well. The sample was rested at room temperature for 5 mins, then centrifuged (11,000 x g, 5 min). The supernatant was collected, isopropanol (200 µL) was added, and the suspension was centrifuged (11,000 x g, 5 min). The supernatant was disposed of, and an aqueous ethanol solution (80%, 1 mL) was added to the pellet, mixed well and centrifuged (11,000 x g, 2 min). Finally, the supernatant was disposed of, the sample was centrifuged briefly, and the remaining liquid was discarded. The pellet was suspended in EB buffer (100 µL). The DNA purity and concentration were determined using a NanoDrop spectrophotometer.

#### 2.2.2 Whole Genome sequencing

High quality extracted DNA samples (1-2 µg) were sequenced at BGI-Australia, QIMR Medical Research Institute, Brisbane, Australia. BGI-Australia constructed DNA libraries for the 17 samples according to BGI-500 in-house library protocols (Kim et al., 2021). The library construction method included the shearing of large genomic DNA fragments into short 200-500 base pair (bp) fragments. Subsequently, purification was performed, and the final products were checked for quality using a Qubit Fluorometer. Later, high-quality DNA fragments were End-Repaired, and A-Tails were added. Following this, adaptor ligation and PCR purification were performed. Finally, the purified high-quality libraries were loaded on the BGI-500 sequencer. All 17 samples were sequenced to high (30X) coverage using BGI-500 with 150 base paired-end sequencing.

### 2.3 Bioinformatics Analyses

Raw reads were obtained from BGI-Australia. The quality of these raw reads was assessed, and adapters were trimmed using the Trimmomatic v3.6 tool (Bolger, Lohse, & Usadel, 2014) with default parameters (ILLUMINACLIP:2:30:10, SLIDINGWINDOW:4:20, MINLEN:50). Trimmomatic was used for BGI reads following a published procedure (Murigneux et al., 2020). The resulting clean paired-end reads were aligned to the human reference genome (hg19) using the BWA v7.3.0 mem module (Li & Durbin, 2009). Subsequently, binary alignment/mapping (BAM) file conversion, sorting, and polymerase chain reaction (PCR) duplicate filtering steps were performed using Samtools v1.3.1 (Li et al., 2009). Next, the Samtools mpileup and bcftools call modules were used to obtain all sites for each sample. Later, multi-allelic (more than two alleles) variants and indels were removed from the VCF files for each sample using an in-house awk script.

For this study, 81 samples that were aligned to hg19 and filtered for PCR duplicate reads (Binary Alignment/Mapping files) were obtained from the Malaspinas et al. (2016) study, including Western Central Desert (WCD; n=13), Cairns (CAI; n=6), Weipa (WPA; n=6), Pilbara (PIL; n=3), Wongatha (WON; n=6), Northern Gold Field (NGA; n=6), Birdsville (BDV; n=6), Riverine (RIV; n=6), Papuan (n=11), African (n=10), European (n=4), and East Asian (n=4) (SI Table 1). All-site genotype calling was performed for each of the 81 samples, and subsequently, indels were removed using an in-house awk script.

Additionally, 12 raw read samples were downloaded from the Mallick et al. (2016) study (https://ftp.sra.ebi.ac.uk/) to increase the sample size for the European (n=6) and East Asian (n=6) populations (SI Table 2). These 12 samples were treated with quality trimming and adapter removal, alignment, and all-site genotype calling as described above.

Finally, all-site VCF files from Aboriginal Australians, such as NPN speakers, PN speakers, Yolngu, WCD, CAI, WPA, PIL, WON, NGA, BDV, and RIV, were merged with worldwide populations, including Highland Papuan (n=10), HGDP-Papuan (n=1), African (n=10), European (n=10), and East Asian (n=10) samples using the bcftools (v1.3.1) software’s merge module (see Abbreviation/Sample ID details in SI Table 1). Next, bi-allelic SNVs were retained from the merged all-site VCF files using an awk script. The merged data comprised 110 samples and 24,865,041 autosomal SNVs, with a 99.01% genotype rate calculated using PLINK (options: --geno).

We also estimated 1st, 2nd, and 3rd-degree relatedness among 15 population samples using PLINK (options: --make-king-table). Furthermore, we calculated the number of heterozygous variants from the 24.86 million SNVs using an in-house script for unadmixed Australian populations such as PIL, WCD, Yolngu, and NPN speakers, along with worldwide populations. Similarly, derived homozygous allele counts were calculated using the ancestral allele from chimpanzees. Additionally, Runs of Homozygosity (RoH) analysis was performed using PLINK (options: --homozyg --geno 0.001 --indep-pairwise 1000 100 0.2) to measure inbreeding and bottleneck effects.

### 2.4 Population genetics analyses

We performed multidimensional scaling (MDS) using the 24.86 million SNVs with PLINK software (Purcell et al., 2007). Admixture analyses were conducted using ADMIXTURE v1.3 software (Alexander, Novembre, & Lange, 2009), with default parameters, except for the (--cv) option, which evaluates cross-validation error (lower values indicate a better fit to the data). This analysis was applied to the SNVs for K values (i.e., number of ancestral populations) between 1 and 8.

For local ancestry analyses, we phased 110 samples without a reference panel using the default parameters of Beagle v4.0 (Browning & Browning, 2007). Local ancestry was further analyzed using RFMix (v2.03) (Maples et al., 2013), incorporating reference populations from European (n=10), East Asian (n=10), and WCD (n=12). We performed masking on 39 samples from six populations with significant European and East Asian admixture, including WPA, CAI, BDV, RIV, NGA, and WON. The RFMix analysis resulted in 4,973,016 SNVs. To maximize the number of SNVs, we retained nine samples from these populations, which exhibited low (10-40%) European and East Asian admixture. Masking of European and East Asian admixture with no missing data yielded 328,395 SNVs.

The final dataset, comprising the admixture-masked samples, included 328,395 SNVs, 42 samples from Australia (SI Table 1), and 40 samples from other global populations for population genetics analysis (SI Table 3). MDS analysis was initially performed using PLINK on the 328,395 SNVs. Following this, we conducted ADMIXTURE v1.3 analysis (Alexander et al., 2009) with the default parameters, except for the (--cv) option, on the same SNVs across K values 1 to 8. We used the VCF2Pop (Subramanian et al., 2019) online tool to process the VCF file containing 82 samples and 328,395 SNVs to calculate genetic distances (i.e., the number of differing SNVs). The VCF2Pop tool generated pairwise diversity data for all 82 samples, which was then used to construct the phylogenetic tree in MEGA (v10) (Kumar et al., 2018).

### 2.5 Statistical analyses

To understand shared genetic drift between populations from within Australia and outside Australia, F3 out-group (Yolngu/A486, pop2; Yoruba) analyses were performed for each sample using admixtool v7.0.1 (Patterson et al., 2012). Here, one of the three populations (Pop3; Yoruba) was considered an out-group and following that, the genetic distance or drift was measured between the remaining two groups. Herein, pop1 (Yolngu) was replaced with one sample, and pop2 was replaced with all the individuals except Africans. F3 analyses were performed before and after masking the ancestry.

### 2.6 mtDNA analyses

Mitochondrial DNA (mtDNA) analyses were carried out using the following samples: Yolngu (n=13) and NPN (n=4) from this study, in addition to 25 mitochondrial genomes from PN speakers, including PIL (n=3), CAI (n=3), WPA (n=2), NGA (n=1), WON (n=1), RIV (n=1), WCD (n=12), and BDV (n=2) from Malaspinas et al. (2016) and Wright et al. (2018). MtDNA sequences were obtained from BAM files of each sample and processed in two steps. First, mitochondrial genomic positions were identified using the Samtools mpileup module. Second, complete mtDNA sequences were generated for each sample using the bcftools consensus module (SI Table 3). Additionally, mitochondrial haplogroups were identified using the HaploGrep 2 (Weissensteiner et al., 2016) web tool, based on the PhyloTree v17 reference database (Van Oven, 2015). The sequences were classified into haplogroups (SI Table 3) using the PhyloTree database. MtDNA multiple sequence alignment analyses were performed using the MUSCLE module in MEGA v10 (Kumar et al., 2018). The alignment data from 42 samples were then used to construct a phylogenetic tree using default (option: Tamura-Nei (TN93) model) in MEGA v10.

## 3 RESULTS

### 3.1 Australian and global genetic diversity

This study sought to understand the genetic relationships among Yolngu speakers from other Aboriginal Australian populations in both a global and an Australian context. Some population size samples used in this study are small (although not for the Yolngu). We acknowledge the limitations of the sometimes-small sample sizes that were available to us. However, it has been argued that, in a whole genome study with millions of markers, two or three samples are sufficient for understanding changes in population genomic history (for example, Rasmussen et al., 2011; Mallick et al., 2016). Nevertheless, we acknowledge that smaller sample sizes limit our ability to fully capture the total genomic variation present in any population, although we believe they are sufficient for our study.

In addition, for this study we selected genomes with no European and / or Papuan admixture and carefully excluded these from the analyses. In summary, these datasets were adequate for the bioinformatic, and evolutionary analyses reported here. We detected a small number of first- and second-degree relatives among the Yolngu, CAI and WCD samples which will have influenced our results, although we believe only to a minor degree.

MDS analyses were performed using 110 samples (Figure 2a). The first two components split the African and non-African populations (including Europeans, East Asians, Aboriginal Australians, and Papuans) because Africans have the highest levels of genetic diversity compared to non-Africans (DeGiorgio et al., 2009). On the other hand, the remaining non-African populations share a common ancestry as part of an Out-of-Africa event. Unadmixed Aboriginal Australians cluster close to Papuans and are more distant from European/East Asian populations (Figure 2a). The MDS analyses of <u>n</u>on-African populations divide them into two distinct groups: Eurasians and Oceanians (including Aboriginal Australian/Papuan individuals) (Figure 2b). Within Aboriginal Australian populations, there are Pilbara and Kalgoorlie individuals within the Western Central Desert cluster and a single Weipa individual in the Yolngu and NPN speakers’ cluster (Figure 2b).

**FIGURE 2.**
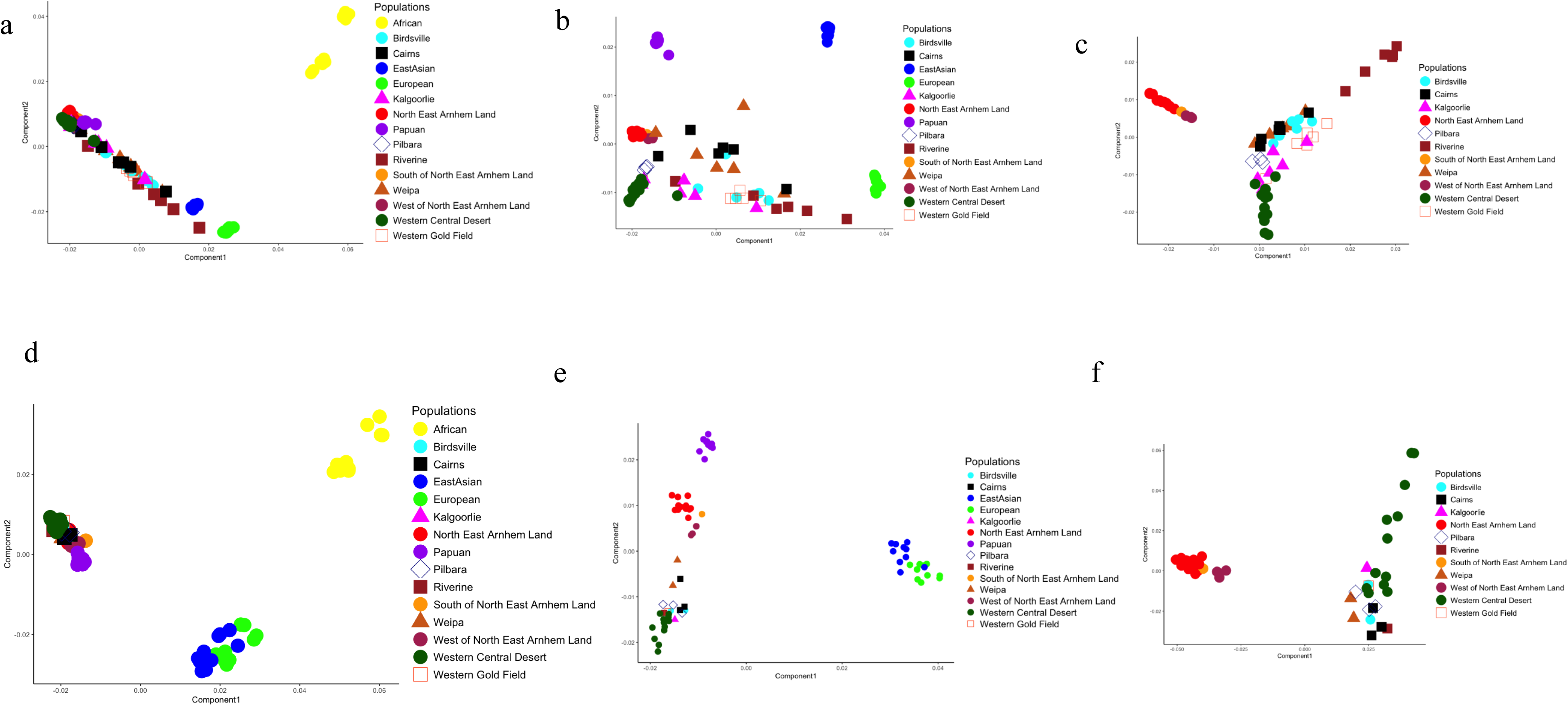
Genetic relatedness analyses of Yolngu speakers and other populations in Australia and worldwide. (a) Multidimensional scaling (MDS) results show genetic affinities before removing Indigenous Australians with European and East Asian admixture, as well as Africans. (b) MDS results show genetic affinities before removing Indigenous Australians with European and East Asian admixture, excluding Africans. (c) MDS plot using only Aboriginal Australians. (d, e, f) plots represent similar data as above but after masking European and East Asian admixture.

The MDS plot also indicates that all the 13 Yolngu people and the 4 NPN speakers whose samples were included in this study have not interbred with Europeans or East Asians, in contrast to samples from other PN-speaking groups such as Cairns, Weipa, Riverine, and Birdsville (Malaspinas et al., 2016; Wright et al., 2018) (Figure 2b). The European and East Asian samples are strongly separated from the Indigenous Australian groups with the Papuans (Figure 2d). After European and East Asian admixture masking, unadmixed samples cluster near WCD and Yolngu genomes (Figure 2d). In addition, the non-African MDS analyses using ancestry masked data form two separate clusters of WCD/ Yolngu and NPN speakers. The remaining Aboriginal Australian samples were spread between the WCD/ Yolngu and NPN speakers, likely explaining shared genetic drift with WCD (Figure 2e). Compared to admixed samples, local ancestry masked Aboriginal Australians form a better grouping (Figure 2c & f). This illustrates the importance of ancestry masking in admixed samples to understand population relatedness.

In addition, we performed analyses of kinship relatedness among the 110 samples used. This revealed that three Yolngu individuals were first- and second-degree relatives (kinship value: < 0.117) while the other 10 samples from Yolngu and four from NPN were unrelated. In addition, two samples from CAI population, and four samples from the WCD population had 1st and 2^nd^ degree relatedness (SI Table 2).

### 3.2 Population Structure

To understand admixture and population structure, admixture analyses were performed before and after removing European and East Asian admixture. For both analyses, K=6 showed the best fit representing six populations used - Africans, Europeans, East Asians, Papuans, NPN, and PN speakers (SI Figure 1a & 1b). Firstly, the admixture plot of 110 samples illustrates the varying level of the six ancestry proportions (Figure 3a). Outside Australia, African, European, East Asian, and Papuan populations had distinct ancestry. Within Australia, Yolngu and NPN speakers’ samples show no European or East Asian admixture, although NPN speakers have PNG and WCD ancestry, whereas the Yolngu do not. The other nine Australian populations show shared ancestry with Yolngu (10-30%) and WCD (20-50%), and most PN samples show recent European and East Asian admixture (10-80%). In addition, the K8 analysis illustrates that NPN and Yolngu speakers had a distinct ancestry (SI Figure 1a). However, more samples from mainland NPN speakers in future studies can shed light on their population structure. Similarly, after removing European and East Asian admixture, samples from Cairns, Weipa, Birdsville, and Riverine show shared ancestry with WCD, Yolngu, and Papuans at a fine scale (Figure 3b). After masking in the admixture plot, Yolngu show population structure. NPN speakers share 7-90% ancestry with Yolngu, 5-10% with Papuans, and 5-10% with WCD. WPA05 showed around 22% shared ancestry with Papuans. Admixed samples from populations from Cairns, Weipa, Birdsville, and Riverine, have no shared ancestry with Europeans or East Asians which suggests ancestry masking removed the admixture. Moreover, WPA02 (12%) and CAI01 (10%) share ancestry with Yolngu and NPN speakers. Admixture plots from K=5 to K=8 show ancestry distribution among samples in the masked samples (SI Figure 1b).

**FIGURE 3.**
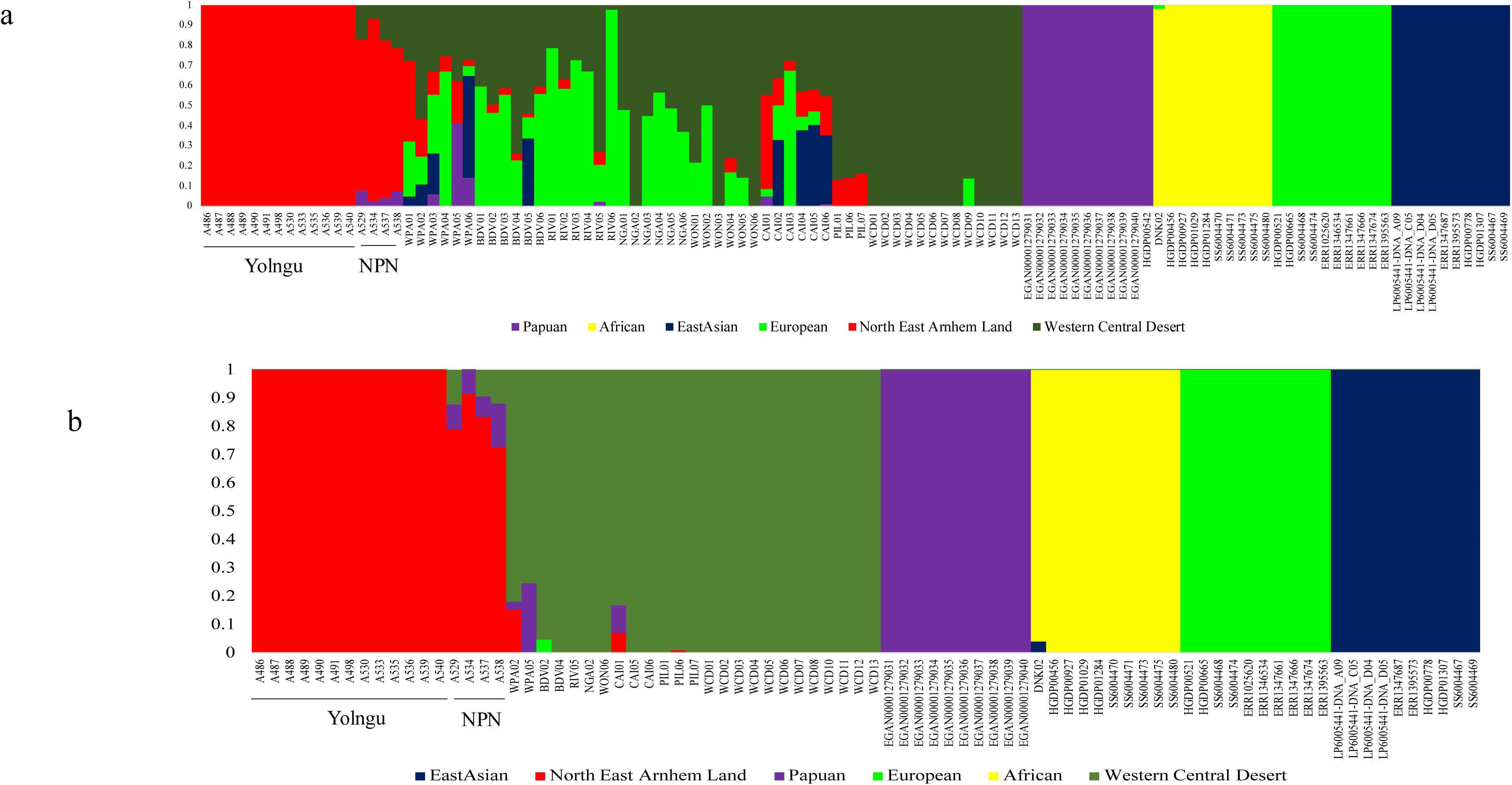
ADMIXTURE plots of Yolngu speakers and other populations showing recent population structure and gene flow among populations before and after removing the European and East Asian admixture. Colours correspond to the collection locations shown in Figure 2. (a) K=6 admixture plot of 110 samples before local ancestry masking; (b) K=6 admixture plot of 82 samples after local ancestry masking.

### 3.3 Genetic relatedness of Yolngu within and outside Australia

To estimate genetic drift and relatedness between Yolngu and the other population samples, f3 statistics were calculated (see SI Table 4). The f3 out-group analyses results show that NPNs (0.229) have the highest mean shared genetic drift with Yolngu people, compared to other populations (Figure 4a). Three Aboriginal Australian populations showed the second highest level of relatedness with PIL, CAI & WCD, followed by Highland Papuan samples. The remaining Aboriginal Australian populations showed low and varying levels of relatedness because of admixture with Europeans/East Asians. East Asians (0.1) showed the second lowest level of relatedness and Europeans (0.09) were the least related to Yolngu speakers. A similar trend was observed in admixture-masked analyses. The NPN were most closely related to Yolngu followed by WPA (0.199) showing the highest mean score compared to other populations such as CAI (0.192334), PIL (0.192006), WCD (0.19311), and BDV (0.185) (Figure 4b). We also carried out genetic drift analyses on all the samples in Yolngu comparing them against other Aboriginal Australian and Papuan populations. The results show NPN speakers have the highest genetic drift values with Yolngu compared to other Aboriginal Australians and Papuans (Figure 4c).

**FIGURE 4.**
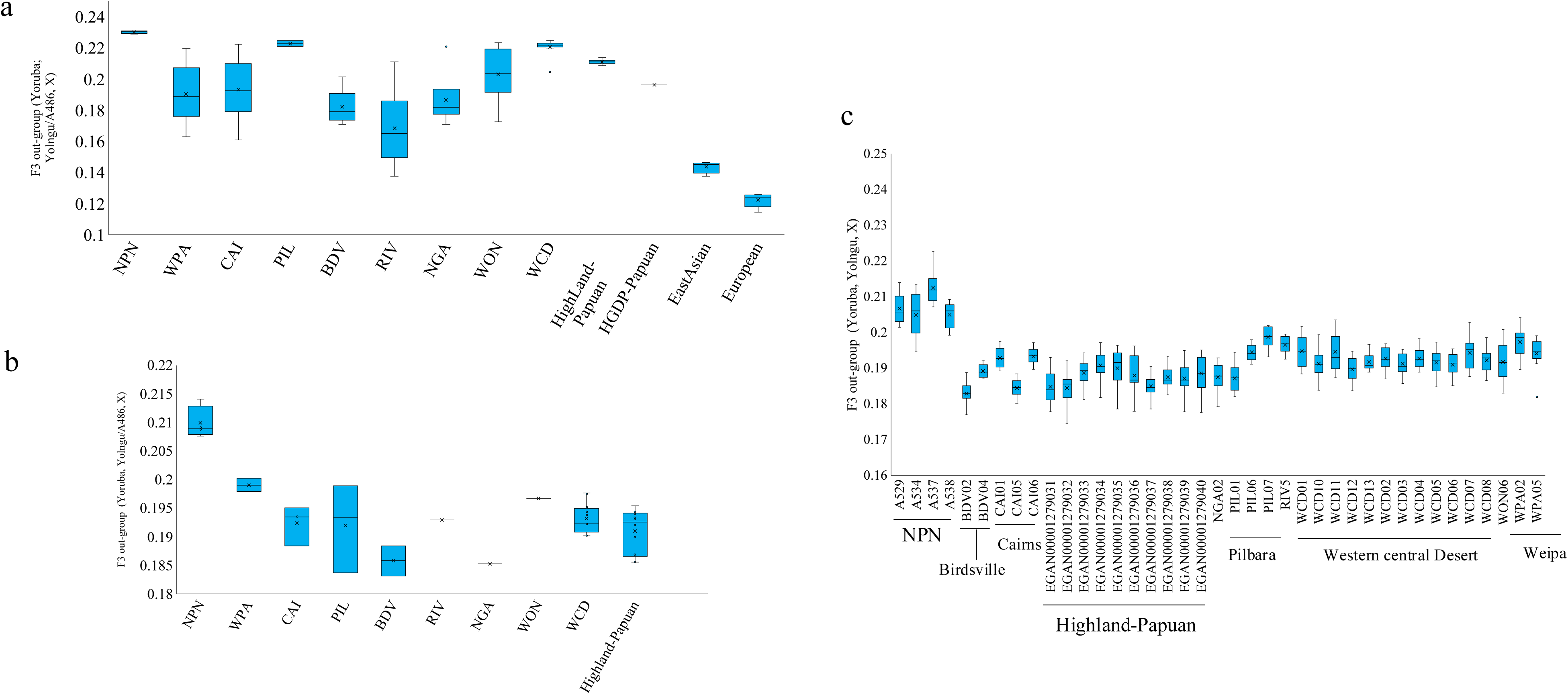
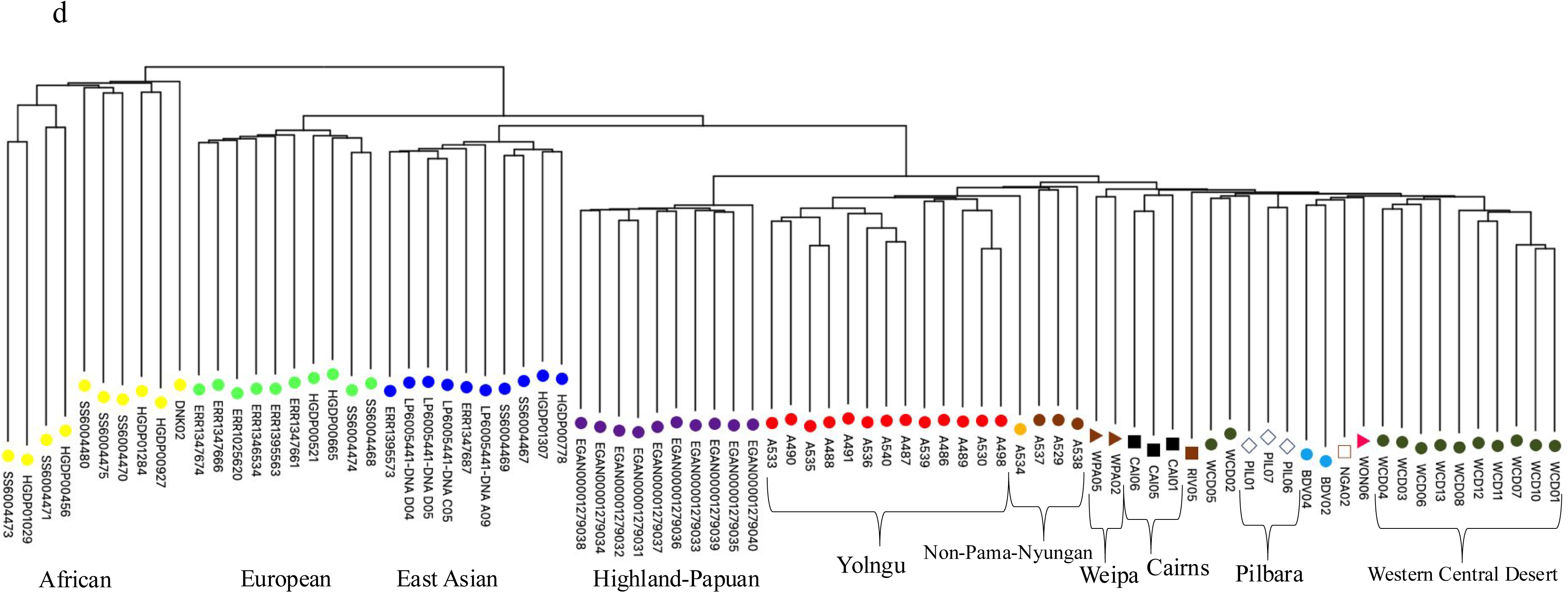
F3 statistics and phylogenetic results. (a) F3 statistics of 13 population samples, including admixed samples, using 24.86 million SNVs. Yolngu show the highest genetic drift with NPN, while populations such as PIL and WCD show the next highest level of relatedness, with Europeans and East Asians showing the lowest relatedness. (b) F3 statistics of population-wise mean counts using one of the Yolngu samples (i.e., A486) show NPN to be the most related to Yolngu, followed by WPA in the ancestry-masked data. CAI, PIL, WCD and Highland Papuans show similar mean shared drift. BDV and NGA have the lowest genetic drift. c) F3 statistics with the Yolngu depict sample-wise variation, which shows that the NPN have the highest relatedness to them, while the remaining samples show similar mean scores. (d) Phylogenetic relatedness among the full nuclear genomes of Yolngu speakers in the context of different Aboriginal Australian PN language groups and four NPN people, after removing individuals with European and East Asian admixture, as well as the genomes from people outside Australia. Colours refer to the collection locations shown in Figures 2a and 2b.

The pair-wise diversity (see SI Table 3) phylogenetic tree results, on ancestry masked data, show Aboriginal Australian relationships with four worldwide populations in a global context (Figure 4d). The Yolngu population (PN) samples were most closely related to the four speakers of the NPN languages, followed by Indigenous people from WPA, CAI, PIL, and WCD. In addition, the NJ tree shows that Papuan samples are an out-group of all the Aboriginal Australian populations including both WCD and Yolngu.

### 3.4 Heterozygosity and RoH

We also measured the heterozygous SNV counts of Yolngu and NPN speakers and compared them to those of other global populations to understand the genetic diversity among populations. First, the mean heterozygous SNV estimates for unadmixed populations show that Africans have the highest level of heterozygosity (2.94 million SNVs), followed by Europeans and East Asians. Among Australian populations, people from the WPA sample (2.13 million SNVs) and CAI sample (2.11 million SNVs) have the highest heterozygosity. The latter had the highest heterozygosity, followed by WCD (1.99 million SNVs), NPN (1.97 million SNVs), and Yolngu (1.92 million SNVs) speakers, who showed the third-lowest levels, just above the PIL population (1.91 million SNVs). Finally, Papuans (1.88 million SNVs) had the lowest levels among all the populations (Figure 5a). The mean homozygosity SNV plot (using derived alleles) shows that Highland-Papuans (2.16 million SNVs) have the highest value. Among Australians, PIL (2.15 million SNVs), Yolngu (2.13 million SNVs), WCD (2.12 million SNVs), NPN (2.09 million SNVs), CAI (2.06 million SNVs), and WPA (2.05 million SNVs) speakers had the next highest homozygous counts, respectively. Africans had the lowest homozygous counts, followed by Europeans and East Asians, suggesting high heterozygosity in African populations (Figure 5b).

**FIGURE 5.**
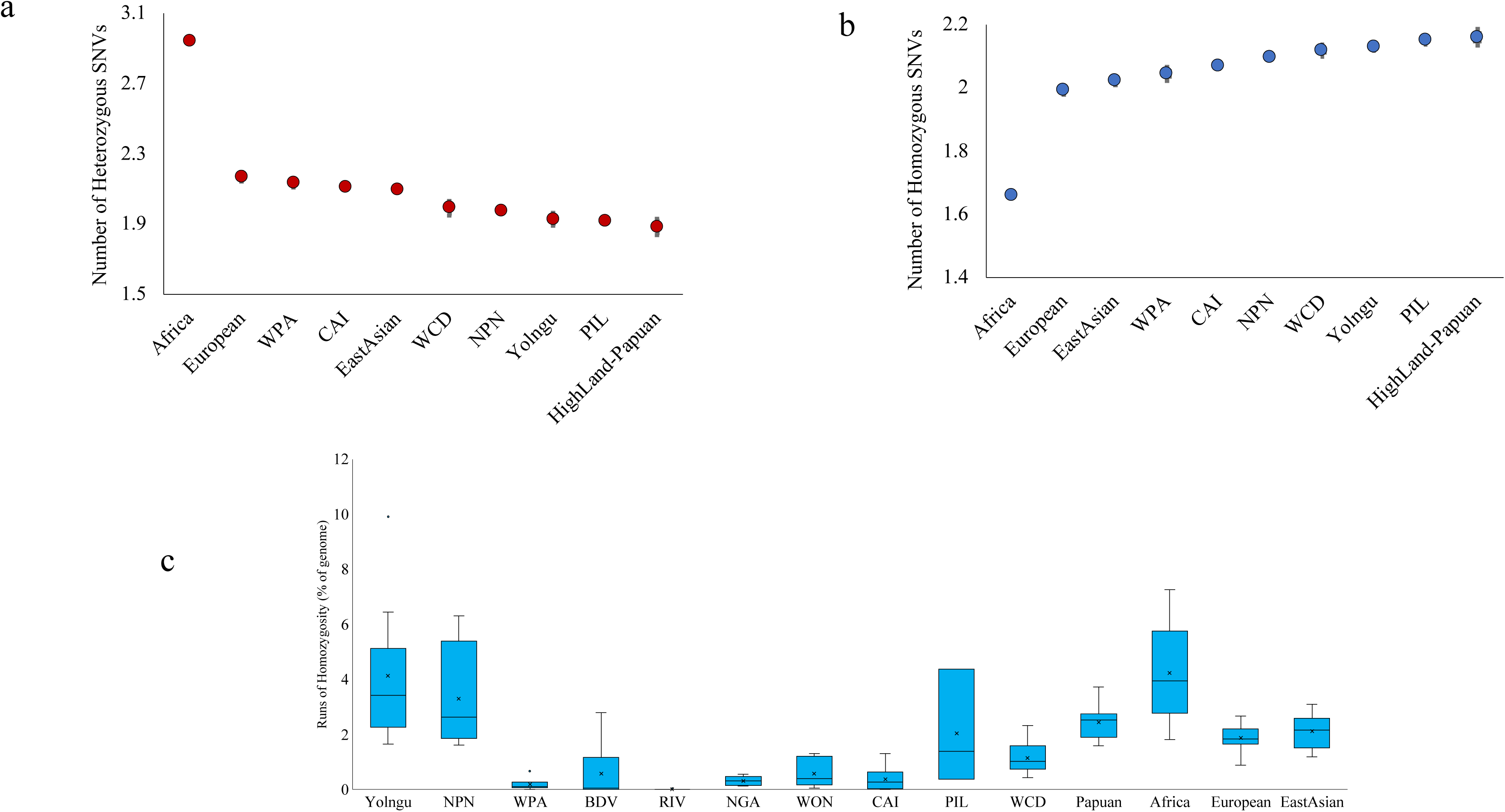
Variation count statistics. a) Mean heterozygous SNV counts for 10 unadmixed population samples using 24.86 million SNVs show that Africans have the highest heterozygosity, while the Highland-Papuan population shows the lowest heterozygosity. b) Mean homozygous SNV counts show that, among the 10 populations, Highland-Papuans display the highest homozygous counts. The Yolngu rank third and NPN speakers have the fifth highest count, suggesting inbreeding or an ancestral bottleneck in the population. (c) Mean RoH in all 110 individuals was compared to assess the level of inbreeding. One Yolngu sample has the highest RoH, representing 10% of the genome.

The RoH length was estimated per individual within each population, considering those greater than 1 Mb in size. Briefly, RoH values can be used to infer inbreeding and bottleneck effects. Yolngu (mean 4.13%) and NPN (mean 3.29%) speakers had the highest mean RoH percentage of the genome compared to other populations, suggesting high inbreeding in Yolngu. Among other Australians, PIL (2.04%) and WCD (1.15%) had greater than 1% mean values, while the others had less than the 1% mean RoH length. Beyond Australia, Africans (mean 4.22%) had the highest mean value (Figure 5c), followed by Papuans (2.44%), East Asians (2.12%) and Europeans (1.87%).

### 3.5 Mitochondrial diversity

For consistency, we used the same number of 42 Aboriginal Australian samples for mtDNA (maternal ancestry) analyses, as used in the nuclear genome analyses after masking. The mtDNA analyses show six major haplogroups, namely N, O, S, P, R, and M (SI Table 3). Yolngu speakers had sub-haplogroups including N/N*, P3a, P5, P7 and S2. In the unrooted phylogenetic tree, N, S and P haplogroups from Yolngu and NPN speakers form two different clades (Figure 6b). The analysis identified a novel N* haplotype among the Yolngu (Figure 6b & SI Table 3). This lineage appears to be young (in comparison to other mitochondrial haplogroups) given the length of the branch, and it shows little diversity (lack of branching structure within the clade). Most of these haplogroups are shared across east-PN, west-PN, and south-PN speakers except for the N haplogroup (Figure 6a). In this study, 55% of N and 45% of P haplogroups were observed in Yolngu speakers (Figure 6b), while the south-west PN speakers including WCD and NGA populations showed 55% M, 15% O, 15% P, and 15% R haplogroup frequencies (Figure 6b). As shown in SI Table 3 and Figure 6b, the Yolngu mtDNA haplogroups (P5, N*, P3a and S2*) differ from the mtDNA haplogroups from the WCD, which are R12, S1a, P3b, O1, O1a, M42 and M14. In addition, S2* appears to be a novel N* haplotype among Yolngu and NPN speakers. They are present in Yolngu (A535) and NPN (A529) individuals.

**FIGURE 6.**
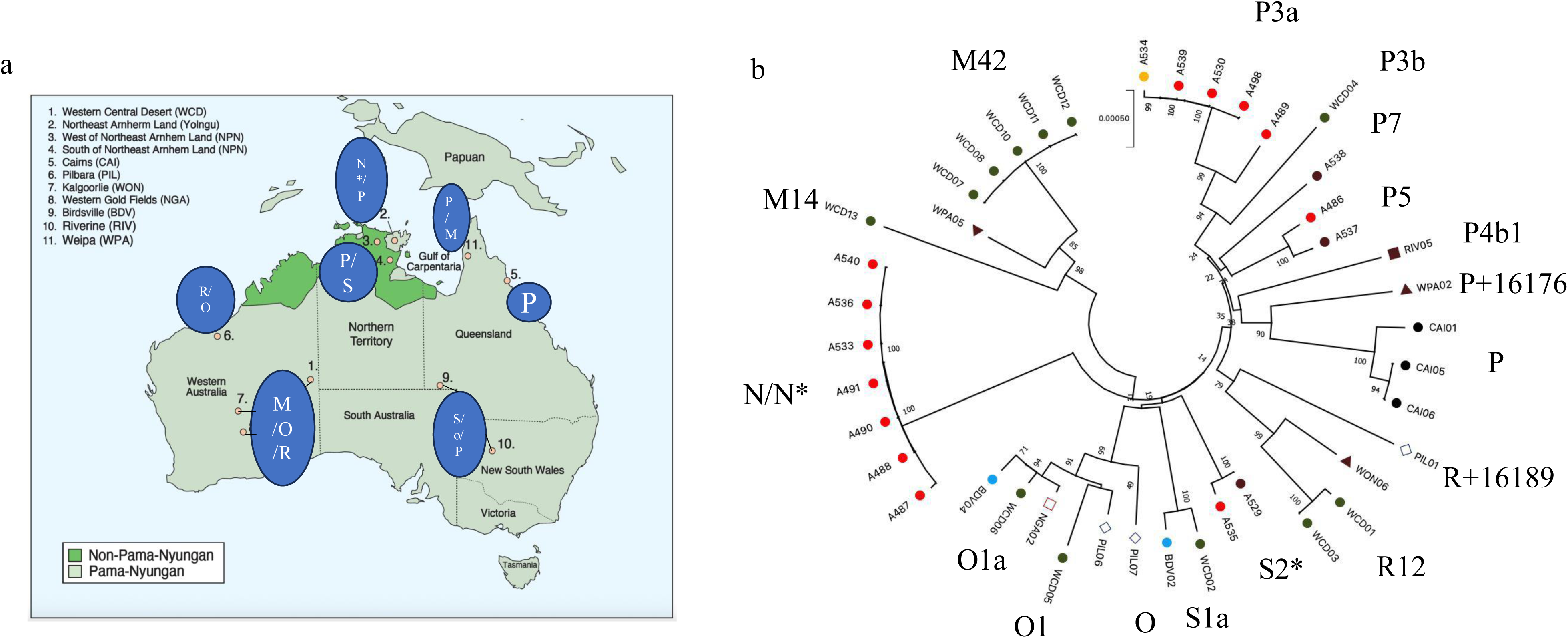
mtDNA phylogenetic tree for Yolngu and otherPN speakers, including four NPN-speaking individuals (see Table 3). Colours correspond to the collection locations shown in Figures 2 and 3. (a) Map showing the distribution of mtDNA haplogroups across regions, identifying six major haplogroups. (b) NJ phylogenetic relatedness diagram showing the distribution of major six haplogroups and subtypes.

## 4 DISCUSSION

Over the last decade, considerable attention has been paid to estimates of genomic variation in Aboriginal Australians. This includes reports of whole genome sequence variation, as well as partial and complete mitochondrial genomes (Malaspinas et al. 2016; Nagle et al., 2017a, b; Tobler et al., 2017; Wright et al., 2018; Reis et al., 2023; Silcocks et al., 2023). Y chromosome variation among Aboriginal Australians has also been investigated over the last decade (e.g. Bergstrom et al., 2016). At the same time, linguists have continued studies on the diversification and dispersal of Indigenous languages of Australia (for example, Bowern 2010, Bouckaert et al., 2018, Evans, 2020, Miceli & Bowern, 2023).

The study presented here is one of the very few and most recent published accounts of the nuclear genomic diversity of Yolngu speakers. Yolngu speakers are important for the understanding of the settlement of Australia owing to their geographical position in the far north of the continent, as they are for an understanding of the spread and diversification of Indigenous languages.

### Yolngu vs WCD Diversity

Figures 2c and 4d reveal a pattern of considerable nuclear genetic diversity among the speakers of the different PN languages, the most marked being between Yolngu and the Western Central Desert and others from the arid interior such as Pilbara populations. This is consistent with previous findings of White (1997) that there is a particularly strong genetic differentiation between these populations based on total ridge counts of fingerprints of Yolngu speakers, compared to other individuals from PN-speaking groups in central Australia. The marked genomic differences within the PN Family, here the Yolngu and the WCD, are in agreement with early genetic studies, showing that the language Family is not a discriminator of genetic identity (White, 1995; 1997). Certainly, the details of genomic diversity within the PN-speaking Family presented here suggest that PN did not spread with the movement of people across the continent; rather, the PN languages diffused among the different populations.

These two regional groups (Yolngu and WCD), which also show strong differences in skin pigmentation (Parsons and White, 1976), come from two markedly different ecological zones. The inhabitants of the WCD and other parts of central and western Australia traditionally occupied large territories in arid to semi-desert areas and had correspondingly low population densities. The Arnhem Land region is characterized by a monsoonal and markedly seasonal climate with considerable habitat diversity, particularly in coastal areas. This diversity provided the Indigenous people with a broader and far more favourable subsistence base. Tribes or language units of this region tend, therefore, to occupy smaller territories and have higher population densities. In addition, marriage distances are far shorter than those in the more arid regions to the south (White, 1979).

The genetic differences shown between the people of central and western Australia and those from eastern Arnhem Land and the Gulf of Carpentaria likely arose as a result of genetic drift and different selection pressures over many generations (Nei, 2013). Such regional differences are consistent with a long period of separation of populations following the north-south movement of people from New Guinea and then across northern Australia. Silcocks et al. (2023) provided estimates of the divergence of Central Australian people (part of the ancestral population shared with Titjikala and Yarrabah populations) from Yolngu (“split times”) of about 31,000 years from nuclear DNA. This is younger than the archaeological evidence, which for the Western Desert is about 50,000 years BP (McDonald et al., 2018). The mitochondrial DNA analysis (SI Table 3; Figure 6) shows that the Yolngu mtDNA haplogroups (P5, N*, P3a and S2*) differ from the mtDNA haplogroups from the WCD, which are R12, S1a, P3b, O1, O1a, M42 and M14. There are no shared haplogroups between the two groups and, therefore, no relationship can be claimed based on the present study. P3a and P3b are distinct from one another and are thought to have separated at least 50,000 years ago. Of course there may have been different arrivals in the centre from different ancestral populations in northern Australia.

### Papuan and Aboriginal Australian relatedness

The analyses presented here show relatedness between Papuans and both PN- and NPN-speaking people (Figures 2 and 4d). The split between Indigenous Australians and Papuans is likely to have occurred about 47,000 to 50,000 years ago (Silcocks et al., 2023; Tobler et al., 2017). Although given an earlier date for the Western Desert, suggested by the archaeological evidence (McDonald et al., 2018; Clarkson et al., 2017), this must be an underestimate. Even though a major colonizing event may have occurred at this time, it is highly likely that before then, and before the sea-level rise and the inundation of the Sahul Shelf about 10,000 years ago, at least small-scale movements of scattered bands of hunters and gatherers occurred over the Sahul Shelf from New Guinea and northern Australia. Presumably there would have been some interaction among them, with limited gene flow. Here we differ from Silcocks et al. (2023), who argue that the ancestral Australian and Papuan populations were genetically isolated by 27,000 to 30,000 years ago. In fact, low-level gene flow has continued across Torres Strait to this day.

### Movement across the Sahul Shelf

Like modern hunters and foragers, these groups would probably have established their own economic ranges and social networks across the bands occupying the area. Could it be that after the area around Lake Carpentaria and the waterways and wetlands across the Sahul Shelf was occupied, people moved through these territories and into the more arid lands to the south, including the Western Central Desert? With an occupation date of about 50 Kya now established for the Western Desert, the spread west around the southern margins of the Sahul Shelf must have occurred at least by this time.

For the Yolngu, dermatoglyphics (fingerprints – total ridge count) and other data based on blood groups, as well as cultural information, led White (1997) to suggest that the Yolngu had their origins to the east of their present location. The nuclear DNA phylogeny (Figure 4) and the MDS plots (Figure 2) show the Yolngu to be most closely related to the NPN speakers who are their immediate neighbours to the west and south of northeast Arnhem Land; and their next closest relationship is with individuals from Weipa and the Cairns area. While the few mitochondrial genomes from these individuals do not share common mitochondrial haplogroups with the Yolngu population, the nuclear DNA phylogeny of the PN speakers studied here does offer support for an ancestral link between PN-speaking people from the Cairns region in the east and those from Weipa on the western side of Cape York, and the Yolngu, with much deeper links to the Papuan people of New Guinea (Figure 2e). However, there are too few mitochondrial samples from Cape York to be definitive. The mitochondrial data are not comprehensive, and the mt lineages do not cluster, while, in contrast, the nuclear data are based on a large number of full genomic sequences, i.e., high coverage biparentally inherited loci. Nevertheless, the small sample sizes for populations east of the Yolngu into Cape York are unlikely to be an adequate representation of those populations. Further genomic studies of these populations are required to test this hypothesis of ancestral links between these groups. The mtDNA analysis identified a novel N* haplotype among the Yolngu. It is yet to be established whether this haplotype also exists in populations to the east of the Yolngu into the western side of Cape York. If further Australia-wide studies of the mt genomes confirm that it is only found among the Yolngu, then it suggests that it either evolved in the Yolngu population in northeast Arnhem Land, or it came with the ancestors of the Yolngu when they colonized their present location, probably from the source of PN in the southern Gulf region. Without age estimates of the lineage, its potential origins must remain highly speculative.

### Yolngu and NPN relatedness

In the sample from northeast Arnhem Land studied here, four NPN speakers were identified (SI Table 3). Of these, three (A529, A537-1 and A538) were immediate western neighbours of the Yolnguand one (A534) was from the southern neighbour of the Yolngu. In the genomic analysis (Figures 2e, 4d and 4b) , the NPN samples cluster with the Yolngu, as was the case with total finger ridge counts (White, 1979), although the admixture analyses (Figure 3) show that the few NPN individuals carry PNG and WCD genomic DNA, which the Yolngu sample lacks, despite there being some low level intermarriage among the groups, at least in recent years. The genetic similarities may extend further west and south. Similarly, the early fingerprint evidence showed no significant difference between Yolngu people and their close neighbours to the east, the Anindilyakwa speakers from Groote Eylandt (White, 1979). Van Egmond and Baker (2020, p. 32) suggested that the Anindilyakwa language

“appears to have undergone separate development for a considerable length of time.” It may be that people occupied Groote Eylandt when it was part of an archipelago and remained on that country after it became an island between 8,000 and 6,000 years ago, although archaeological evidence for this is yet to be identified.

### Yolngu and PN relatedness

The study presented here provides additional knowledge regarding the origin of Yolngu speakers, their settlement in the northeast of Arnhem Land and their genetic relatedness to other PN-speaking populations across Australia. Nuclear genomic analyses suggest that Yolngu have diverged from populations on the western side of Cape York, such as those from around Weipa (Figures 2e and 4), before the Holocene. Previous fingerprint analyses also indicate that there is a close genetic relationship between the Yolngu and people from other populations (both PN and NPN) from around the Gulf of Carpentaria to the east, through to Mornington Island (White, 1979).

After divergence of ancestors of the Yolngu from groups in Cape York, before the Holocene, the migration of early Yolngu from the east may well have occurred during the early Holocene period of increasing sea levels which caused people to move (Smith et al., 2011; Bradshaw et al., 2021). At 11-8 Kya, the Sahul Shelf flooded Lake Carpentaria, although Mornington Island, Groote Eylandt, the Wessel Islands, and the Tiwi Islands were still attached (Williams et al., 2018). Before that time, the Sahul Shelf was in place with Lake Carpentaria, while the coastline of northeast Arnhem Land was about 170 km further north from Buckingham Bay. Between 8-6 Kya, these islands were formed as the sea levels rose to their present height.

This movement of ancestral Yolngu may have coincided with a possible PN language expansion about ∼7,000 years ago (Bouckaert et al., 2018; Evans & Jones, 1997), during which these people and/or their language could have moved from the east around Lake Carpentaria into Arnhem Land, although this date must remain a loose estimate. The Yolngu may have travelled through the source area of the PN languages somewhere in the southern Gulf when they acquired their PN language. These ancestral Yolngu PN language speakers, or the languages themselves, may then have spread through the existing NPN-speaking groups into their present location. (The age of NPN languages and their presumed evolution from proto-Australian is not known.) Alternatively, early Yolngu were in place in northeast Arnhem Land speaking proto-Australian or early NPN languages before their PN language was adopted. This might be the reason for the rapid decline in effective population size and the increase in population structure for the Yolngu, reported by Silcocks et al. (2023). They claim their research provided strong support for an extended period of large effective population sizes before undergoing a structure reduction at about 6,000 years ago, which was followed by a period with small but stable population sizes. Although not considered by Silcocks et al. (2023), this could have been the result of the spread of PN language(s) into Yolngu country with the establishment of language and other socio-cultural boundaries. These boundaries would have restricted gene flow between language groups and communities, leading to an increase in population structure within the Yolngu and surrounding groups, just as has been found for contemporary tradition-orientated Yolngu (White, 1995). NPN speakers of the Gunwinyguan languages (a possible branch of a large language Family from Arnhem Land) may have moved west and established the southern boundary of the Yolngu. It is intriguing that the four NPN-speaking individuals in this study, but none of the Yolngu (also noted by Silcocks et al., 2023), showed Papuan admixture (Figure 3b). This is consistent with the early Yolngu moving west from an area not occupied by people with genetic links to earlier populations in New Guinea, or that this was a founder effect.

Early fingerprint studies (White, 1979, 1997) are informative in exploring population genetic relationships and genetic structure. This was particularly so before genomic techniques became available. Confirmation of their value is shown in this study with the differentiation of people in central Australia from Yolngu in northeast Arnhem Land shown by fingerprints and the more recent genomic studies, such as reported here and by Silcocks et al. (2023). Silcocks et al. and Reis et al. (2023) found the Tiwi people to be a genetic out-group, just as was found by fingerprint analyses (e.g., White, 1979). This provides confidence in the use of dermatoglyphs in exploring genetic relationships between the Yolngu and groups to the east. These data show that the PN-speaking people of northeast Arnhem Land, the Yanyuwa of the southwest Gulf coast and the NPN groups from the Gulf of Carpentaria are closely related, more likely through common ancestry. That is, the linguistic diversity in this region has occurred within a set of genetically related populations. Alternatively, it is plausible that at least a small number of PN-speaking people spread into the northeast Arnhem Land enclave, as a result of displacement around the Gulf of Carpentaria, or for trade and exchange and/or for wives, taking the language with them into a proto or full NPN population. The migration of people from the east around the southern Gulf into northeast Arnhem Land, as suggested by the genetic evidence, receives some intriguing support from Yolngu religious / spiritual beliefs. Aspects of Yolngu mythology are eastward looking (White, 1997). While research by Silcocks et al. (2023) indirectly supports this hypothesis by calculating the timing of split events, preliminary studies by this group have shown the need for more detailed studies to identify precise divergence times. This team aims to estimate such divergence times in a future study including other populations from a previous study (Kumar, 2021).

Further genomic studies of populations to the east of the Yolngu into Cape York are required to test this hypothesis. Consistent with this suggested movement and links to the east is the research on the internal phylogeny of members of the PN language Family (Bowern & Atkinson, 2012), which closely groups Yolngu with the Warluwaric language from the PN language Family on the Queensland–Northern Territory border. PN languages also appear to have spread from this northern part of Australia into genetically different populations to the south, including the WCD. In this case, it seems clear that the languages dispersed and not the people. This supports the conclusion of the few other scholars (summarized by Bouckaert et al. 2018) who also point to the spread of PN languages without the replacement of genes. There is no evidence that the different language Families have genetic origins beyond Sahul.

In summary, Figure 4d shows that there is a strong monophyletic relationship between the nuclear genomes of Yolngu samples studied here, separate from those from the WCD. The nuclear genome data presented here indicate that the most closely related outgroups to this clade likely include PN-speaking populations from Cairns on the east coast and those from Weipa in western Cape York (Figure 4). This genetic relationship between Yolngu and PN speakers from Cape York, which is likely to have been established before the Holocene, hints at the later movement of people around Lake Carpentaria during the Holocene which resulted in the establishment of the Yolngu people and their language (Figure 1).

### Future directions

Future studies of the genomes of NPN speakers from northern Australia will extend our understanding of the evolution of both PN and NPN speakers as well as our wider understanding of the relationship between genome and language evolution. This study points to a more complex process of settlement of Aboriginal Australia than has been demonstrated previously in studies of the genomes of PN-speaking groups south of the area of linguistic complexity in the north (Malaspinas et al., 2016). The research presented here is focused on the relationships and divergence of populations who now speak either PN or NPN languages. However, care must be taken in using genomics (or dermatoglyphs) to elucidate the history of the languages themselves. It is essential to stress that genomics relate to the people and not their languages.

We examined two hypotheses; firstly, that the PN language Family spread among other populations, and secondly, that people speaking a PN language moved into other areas, taking the language with them. The fact that the Yolngu and the WCD, and other PN-speaking groups that are included in this study, differ genomically from each other, supports the first hypothesis. To advance the genomic studies and the understanding of the arrival and spread of Australia’s first peoples, it is necessary to analyse the genomes of NPN-speaking groups both to the east and west of northeast Arnhem Land across to the Kimberley. It is still to be determined if the ancestors of the Yolngu in northeast Arnhem Land moved into the area from the east around the gulf of Carpentaria, speaking a PN language, or whether the PN language spread through NPN-speaking neighbours into an earlier proto-Yolngu population who may in fact have spoken a NPN language, or whether indeed these neighbours moved around the south of the Yolngu and hence separated them from other PN speakers to the south and southeast.

## AUTHOR CONTRIBUTIONS

Neville White’s long relationship with and understanding of the Yolngu people facilitated this research. All authors wrote the original manuscript. Manoharan Kumar performed the genomic DNA extraction and data analyses.

## ACKNOWLEDGMENTS

We thank members of the Yolngu community of Gapuwiyak for their permission to conduct this study and for their approval of this publication, in particular Ms J.Y. Guyula for her great help with the collection of specimens and support for the project.

We appreciate the funding support of the Australian Research Council to David Lambert. We thank Dr. Uma Ramasamy from the University of the Sunshine Coast and Prof Craig Millar from the University of Auckland for their collaboration over a long period. We appreciate the assistance of Vivian Ward also from the University of Auckland, for her assistance with graphics.

## DATA AVAILABILITY STATEMENT

Griffith University Human Ethics approval (GU ref no: 2022 / 207) was granted to David Lambert, the study data are currently being uploaded to the EGA European Genome-Phenome Archive and will be available based on EGA Data Access Committee approval policy.

