## Supplementary Figures for "A high-resolution genomic study of the Pama-Nyungan speaking Yolngu people of northeast Arnhem Land, Australia"

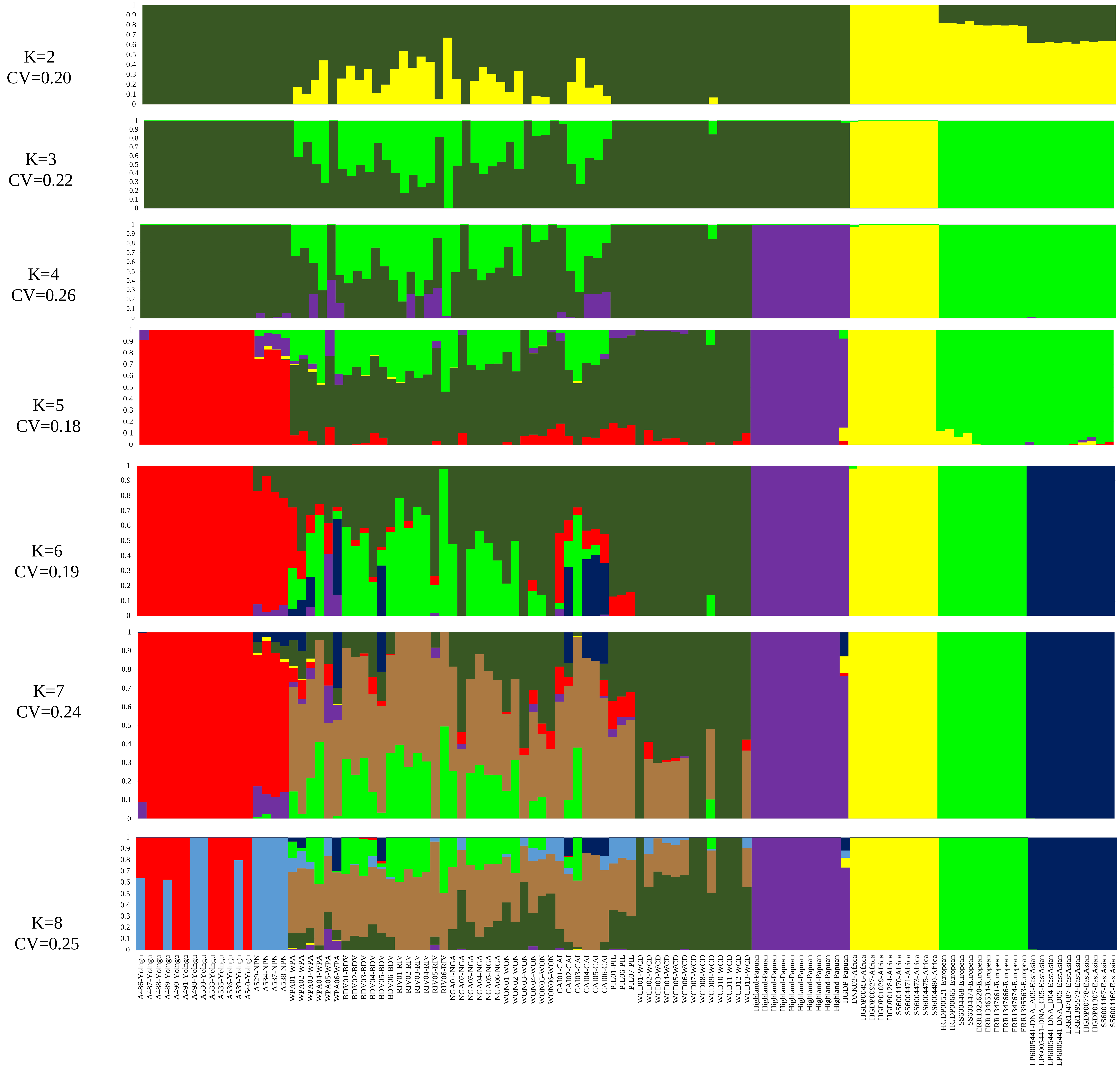


FIGURE 1a Admixture plots between K=2 and K=8 illustrate ancestry proportions for the 110 samples. In addition, CV error score showed the lowest score at K=5. In these admixture plots Yolngu and NPN speakers show population structure at K=5.

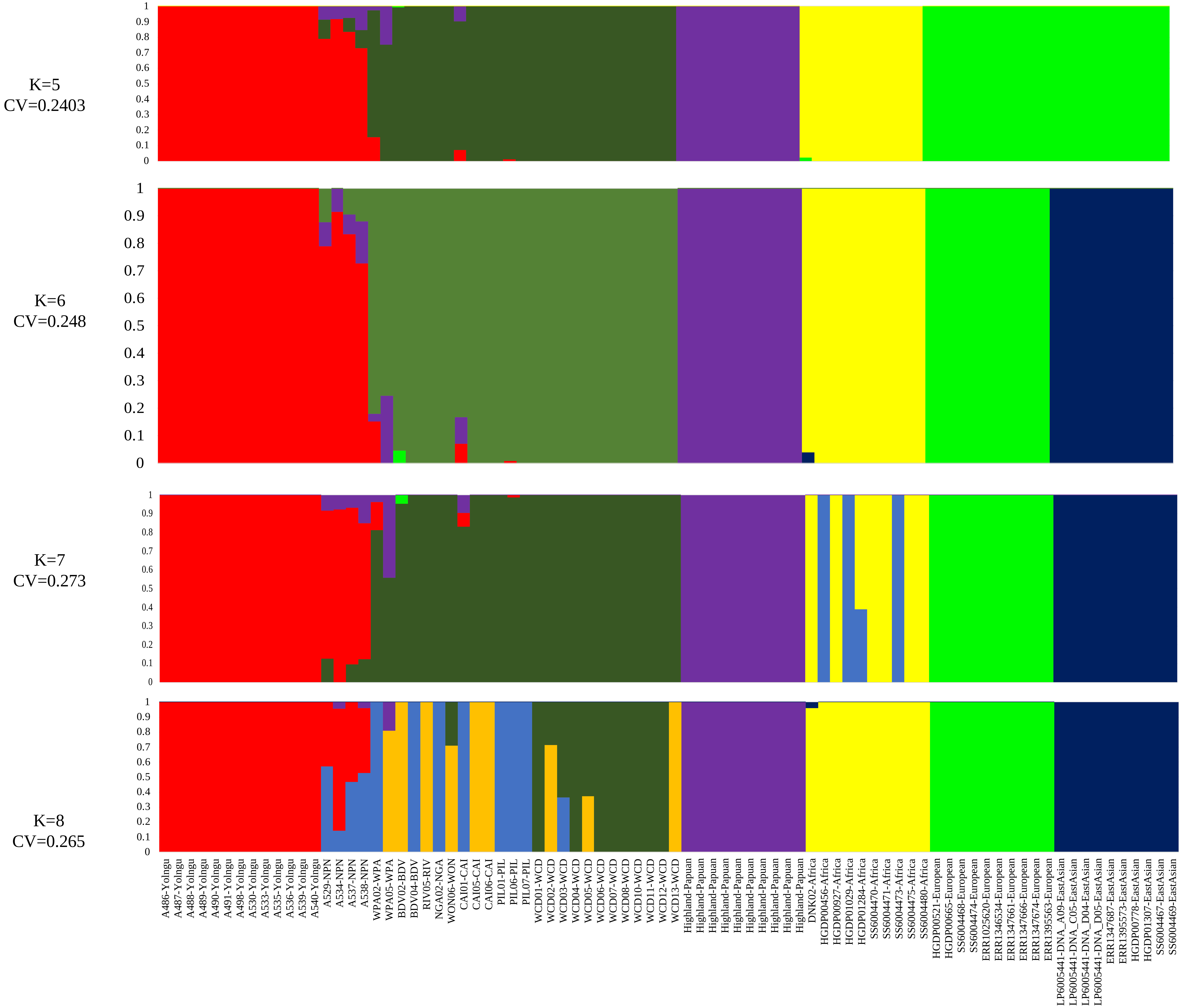


FIGURE 1b Ancestry masked admixture plots between K=5 and K=8 illustrate 82 samples ancestry proportions for 82 samples. The, CV error score showed the lowest score at K=5, illustrating population structure for Yolngu and NPN speakers.
